# A novel VPS33A interaction motif is a master-switch controlling the inhibition and promotion of mammalian Stx17 interactions with SNAP29 and VAMP7 during autophagy

**DOI:** 10.1101/182626

**Authors:** Rebecca S. Saleeb, Deirdre M. Kavanagh, Alison R. Dun, Paul A Dalgarno, Rory R Duncan

## Abstract

Autophagosome clearance is accomplished by SNARE-mediated fusion of the vesicle membrane with the endolysosome. This must be carefully regulated to maintain the organisation of the membrane system and prevent mistargeted degradation. Here, we dissect the autophagosomal SNARE complex pathway and its regulation using FLIM-FRET as a readout of protein interaction *in situ*. We show that within the spatio-temporal framework of the cell, autophagosomal Stx17 preferentially heterodimerises with SNAP29, subsequently associating with VAMP7, not VAMP8 as currently believed. Additionally, we identify for the first time multi-modal regulation of SNARE assembly by the SM protein VPS33A, finding parallels and differences with other syntaxin-SM interactions and suggesting a unifying model of SM regulation. Contrary to current theory, the Stx17 N-peptide interacts in a positionally conserved, but mechanistically divergent manner with VPS33A, providing a late ‘go, no-go’ step for autophagic fusion *via* a phosphoserine master-switch.

## Introduction

Macroautophagy (henceforth autophagy) is a bulk intracellular degradation pathway that transports cytoplasmic material to the lysosome for hydrolysis. Autophagy has an important role in protein turnover, the elimination of cytotoxic material and energy homeostasis. The wide range of conditions linked to autophagy^1^ highlight its importance in health and disease.

Autophagy sequesters cargo by the growth of an open-ended double-membrane vesicle, named a phagophore, which forms *de novo*^2^ upon nucleation at ER-mitochondrial contact sites^3^. Closure of this structure forms an autophagosome, which is transported to the endolysosome where it deposits its contents by membrane fusion. This isolation of cargo by sequestration rather than vesicle budding is a seemingly unique mode of membrane trafficking.

Soluble N-ethylmaleimide sensitive factor attachment protein receptor (SNARE) proteins are established molecular drivers of membrane fusion^4^. A subset of these proteins form a fusion event-specific four alpha-helical complex, composed of a Qa-, Qb-, Qc- and R-SNARE motif^5^, which spans the apposing membranes. Amalgamation of the lipid bilayers is promoted by ‘zippering’ of the SNARE complex from the N- to the C-terminal^6^.

Despite the apparently unusual form of membrane trafficking in autophagy, autophagosome clearance is a SNARE-mediated process^7^, suggesting parallels may exist with the well-studied membrane fusion event of regulated exocytosis. Little is known, however, about the acquisition and regulation of the fusion machinery involved in autophagosome clearance, which together determine fusion competence, membrane specificity and spatio-temporal control.

Syntaxin 17 (Stx17), which decorates autophagosomes, has previously been identified as the resident autophagosomal Qa-SNARE^7^. Mammalian Stx17 has been shown to associate with the soluble Qbc-SNARE, SNAP29, and the endolysosomal R-SNARE, VAMP8, *in vitro*^7^. However, a functional interaction has never been reported between these SNAREs *in situ*. Fluorescence colocalisation is commonly used to confirm *in vitro* interactions, yet, diffraction-limited microscopy is restricted to ˜250 nm resolution^8^, from which molecular association cannot be concluded. This distinction is particularly relevant to studies of autophagosome clearance; the convergence of autophagy with other trafficking pathways could lead to the non-functional accumulation of SNAREs on the lysosomal membrane, while the binding promiscuity of SNARE proteins^9^ may affect *in vitro* data.

Organisation of any membrane system is dependent on careful regulation of fusion specificity, for which combinatorial SNARE complex formation alone is not sufficient^10^. Regulation may be accomplished by Sec1/Munc18 (SM) proteins, which have been observed to have multiple regulatory mechanisms, including both the promotion and inhibition of fusion, dependent on the SM-SNARE pair and their binding mode^11^. Fusion promotion is accomplished by stabilisation of the SNARE bundle^12^, 13, the enhancement of complex fusogenicity^14^ or the recruitment of SNARE proteins to the fusion site^15^, with inhibition mediated by stabilisation of a ‘closed’ conformation of syntaxin^16, 17^.

The SM protein VPS33A reportedly promotes both SNARE-mediated autophagosome clearance^18^ and late endosome-lysosome fusion^19^. VPS33A is unique among mammalian SM proteins in forming an integral part of multi-subunit tethering complexes, one of which is the late endosomal HOPS (homotypic fusion and vacuole protein sorting) complex^20^, from which it likely modulates autophagosome clearance^21^. Importantly, VPS33A is also divergent in being considered as an SM protein that cannot interact with a cognate syntaxin N-peptide region, because it lacks an acceptor binding pocket thought to be essential for such an interaction mode^22^.

In the present study, autophagosomal SNARE interactions were studied *in situ* using fluorescence lifetime imaging microscopy (FLIM) to detect Förster resonance energy transfer (FRET)^23^. With this technique, we have investigated the regulation and formation of the autophagosomal SNARE complex in cells. Our data show that contrary to current models, VAMP7, not VAMP8, is the dominant endolysosomal R-SNARE. Additionally, the clear punctate binary SNARE heterodimer pattern we identify highlights an as yet uncovered mechanism for the negative spatio-temporal control of Stx17 interactions elsewhere that must exist in order to prevent ectopic fusion. We demonstrate here that Stx17 fusion-competency is regulated by an N-peptide phosphosite, presenting a hitherto unidentified regulatory step in autophagy. We show that its phosphorylation status mediates a novel second mode interaction of VPS33A, contradicting its accepted classification as a non-inhibitory SM protein^22^.

## Results

### Stx17-resident autophagosomes co-occur with SNAP29 and VAMP7

Stx17 is well established as the autophagosomal SNARE^7^, redistributing to punctate structures colocal with the autophagosome marker, LC3^24^ upon the induction of autophagy (supplementary figure 1). To verify that the observed EGFP-Stx17 puncta in autophagic HeLa cells are autophagosomal vesicles, we used super-resolution microscopy to detect the membrane targeting of Stx17 in LC3-positive puncta (figure 1a-b). Our continuous-wave gated stimulated emission depletion (CW-gSTED) microscope can resolve to 50 nm (supplementary fig 2), providing five-fold improvement on diffraction-limited confocal laser scanning microscopy (CLSM). As expected for a correctly folded integral membrane protein, EGFP-Stx17 is restricted to LC3-resident ring structures when super-resolved (figure 1a-b). Additionally, the correlation between mCherry-Stx17 and EGFPLC3, which is hydrolysed following fusion^25^, increases when fusion is chemically blocked with bafilomycin A_1_ (supplementary figure 1), suggesting that these are indeed fusion-competent autophagosomes.

**Figure 1.**
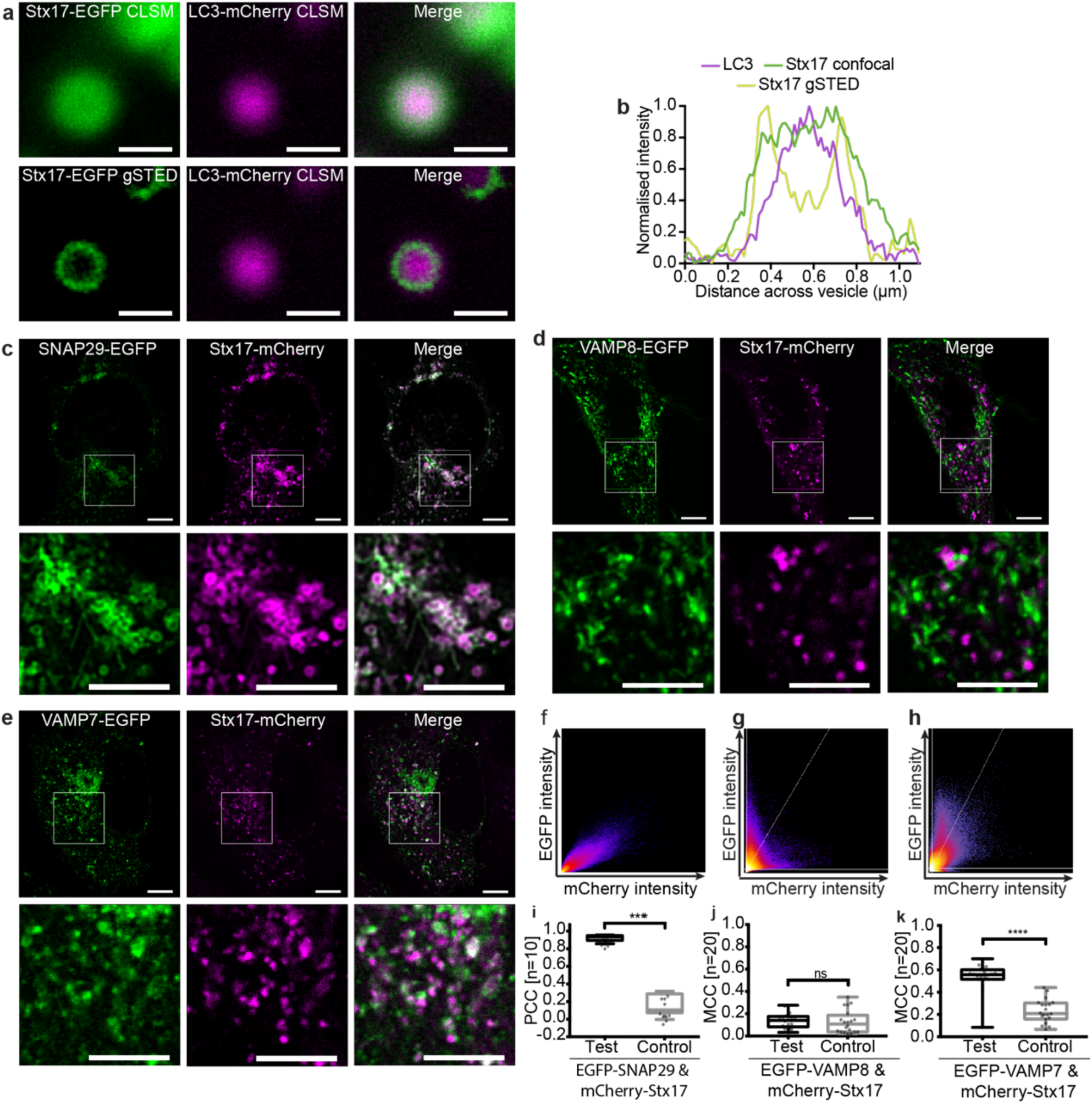
Autophagosomal syntaxin17 colocalises significantly with SNAP29 and VAMP7. (**a**) Comparative confocal and gSTED images of membrane-bound Stx17 rings (left) surrounding LC3-labelled autophagosomes (centre) and a merge of these images (right) from rapamycin treated HeLa cells, scale bars 500 nm. (**b**) Intensity line profile through the punctum in (a) demonstrating membrane localisation of Stx17 resolved by gSTED. (**c-e**) Single channel and merged images of rapamycin-treated HeLa cells expressing Stx17 with either (**c**) SNAP29, (**d**) VAMP8 or (**e**) VAMP7 and a merge of both channels. Scale bars are 10 µm for full field and 3 µm for zoomed regions. (**f-h**) Frequency scatter plots of reported single pixel intensity values in each channel presented in (c-e). (**i**) Quantification of SNAP29 and Stx17 colocalisation using Pearson’s correlation coefficient (PCC) demonstrating significantly higher correlation than in negative controls generated by single-channel 90° rotation. (**j-k**) Quantification of Stx17 puncta colocal with VAMP8 (**j**) or VAMP7 (**k**) puncta using Manders correlation coefficient. All box-and-whisker plots represent the median (central line), 25^th^ and 75^th^ quartile (box) and the minimum and maximum value (whiskers).

Fluorescence colocalisation studies have been used here to verify the expected signal correlation between SNAREs in autophagic cells using CLSM with experimentally verified channel alignment (supplementary figure 3). Stx17 and SNAP29 demonstrate strong signal correlation that follows a linear relationship and significantly co-varies as determined by Pearson’s analysis (figure 1c, f and i). Perhaps surprisingly, however, poor signal correlation is evident with VAMP8 (figure 1d, g and j). An alternative endolysosome-resident R-SNARE, VAMP7, instead demonstrated partial, strong, correlation with Stx17 (figure 1e, h and k), as would be expected if their co-targeting is restricted to post-fusion compartments. This hints at a multi-step assembly process analogous to the regulated exocytotic complex^26^, where the Q-SNARE binary heterodimer first forms, incorporating the R-SNARE in a short-lived *trans*-SNARE complex that spans apposing membranes, becoming a *cis*-complex following fusion. The colocalisation of such sub-populations are poorly suited to metrics of co-variance (supplementary figure 3) and instead the fraction of signal co-distribution was ascertained by Manders correlation coefficient^27^, showing significant correlation of Stx17 with VAMP7, but not VAMP8 (figure 1j-k). Importantly, unlike VAMP8, we find that VAMP7 also correlates with LC3 in a fusion-dependent manner (supplementary figure 4).

In contrast to current thinking about mammalian autophagy, the R-SNARE governing autophagosome fusion in *Drosophila melanogaster* is VAMP7^28^. This discrepancy with the proposed mammalian complex is broadly accepted because *D. melanogaster* has no equivalent to VAMP8; VAMP7, which is 62% similar (supplementary figure 5), is considered its closest homolog. However, unlike VAMP8, VAMP7 encodes a regulatory longin domain, also present in *D. melanogaster* VAMP7 and the *S. cerevisiae* autophagosomal R-SNARE, Ykt6 (supplementary figure 5). This is suggestive of a conserved functional role that highlights VAMP8 as a curious candidate SNARE for autophagy.

Furthermore, homotypic and heterotypic endosomal fusion events appear to be differentially mediated by VAMP8 and VAMP7^29^, respectively, placing VAMP7 as an arguably more credible candidate for the heterotypic fusion of the autophagosome and endolysosome. Of course, as discussed previously, colocalisation does not mean interaction, necessitating more sophisticated approaches to discriminate complex formation from fluorescence overlap.

### SNAP29 and VAMP7 interact with Stx17 *via* its SNARE domain

Protein interactions can be inferred using FLIM-FRET, enabling us to verify, *in situ*, the autophagosomal SNARE complex. As illustrated by the cartoon in figure 2a, FRET is the non-radiative transfer of energy from a donor fluorophore (EGFP) to an acceptor (mCherry), requiring sustained proximity of the fluorophore pair, typically driven by protein interaction. The occurrence of FRET causes the donor fluorescence to decay more rapidly; quantification of this property as 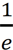, termed its fluorescence lifetime, provides a robust metric for proximity. As protein proximity increases, the FRET efficiency increases, causing fluorescence lifetime to decrease. For the FRET pair EGFP and mCherry, 50% FRET efficiency theoretically occurs at 5.24 nm separation^30^ and requires sub 8 nm proximities for FRET to occur, evidenced by FRET efficiencies of 0.1 and greater (figure 2b).

**Figure 2.**
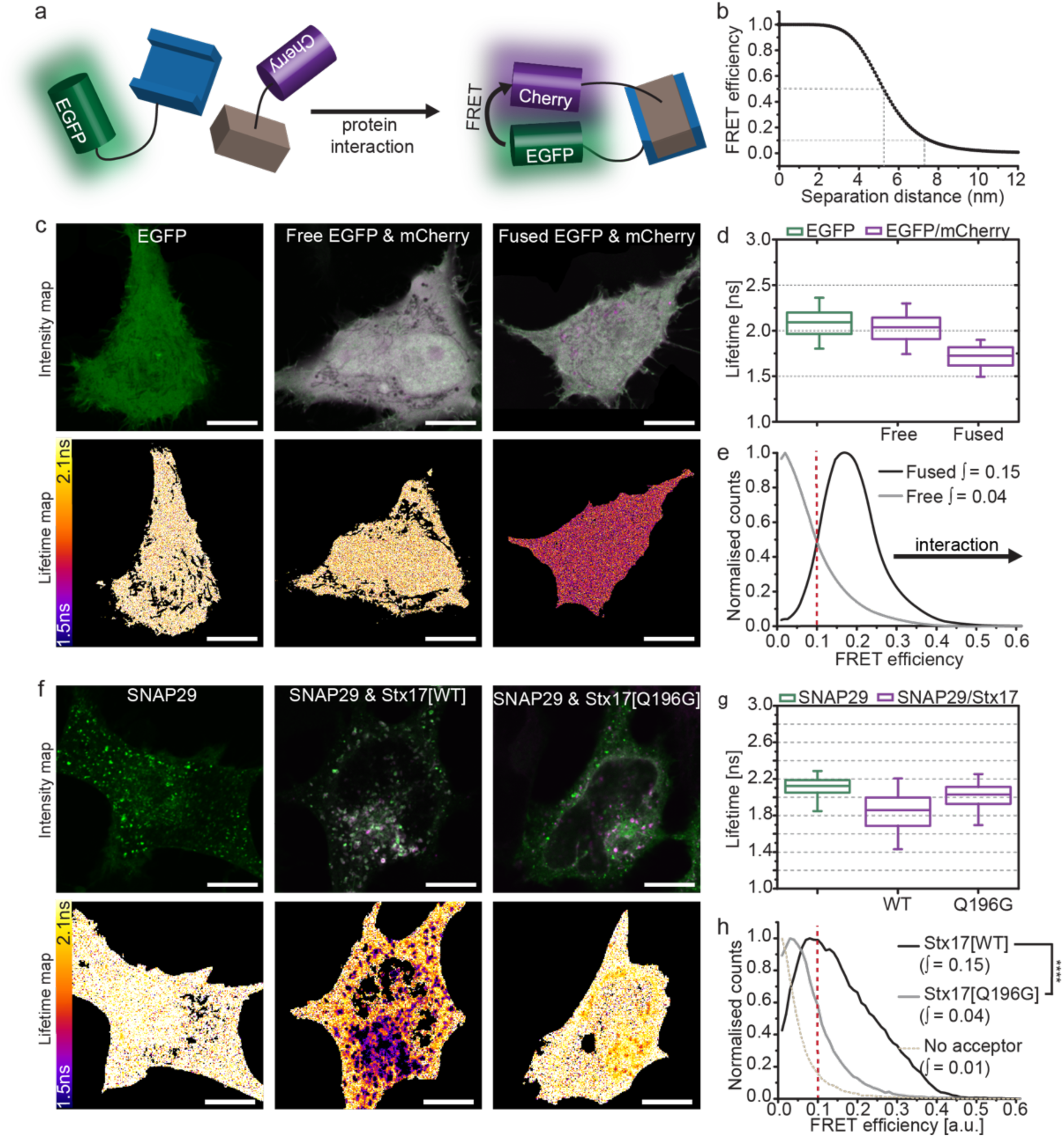
FLIM-FRET confirms Stx17 and SNAP29 interact in autophagosomes. (**a**) FLIM-FRET can be used to probe protein interactions *in situ*. Upon protein interaction, the fluorophores are held in sustained proximity, allowing FRET to occur between donor (EGFP) and acceptor (mCherry) resulting in a reduced lifetime of the donor and acceptor stimulated emission. (**b**) The theoretical relationship between FRET efficiency and separation distance for EGFP and mCherry demonstrating the sensitivity of FRET efficiencies above 0.1 to protein proximities less than 8 nm. (**c**) Intensity and fluorescence lifetime images of unfused, soluble EGFP, co-expressed unfused-EGFP and -mCherry or directly fused EGFP-mCherry. (**d**) Fluorescence lifetime boxplots derived from single pixel analysis of the data set represented in (c). (**e**) Histogram of the normalised single-pixel FRET efficiency counts for free or fused EGFP-mCherry (accumulated from four cells per sample). Integrated values above 0.1 are given, protein proximity causes a right-shift in FRET efficiency and an increase in the integral. (**f**) Intensity and lifetime maps of EGFP-SNAP29 alone or co-expressed with either Stx17[WT] or Stx17[Q196G] in rapamycin-treated HeLa cells. (**g**) Boxplots of all single pixel fluorescence lifetimes accumulated across four cells per condition in (f) and (**h**) the equivalent histograms of normalised single-pixel FRET efficiencies. Integrated values above 0.1 were tested for statistical significance using a one-tailed unpaired two-sample t-test [n=4]. Box-and-whisker plots represent the median (central line), 25^th^ and 75^th^ quartile (box) and 1^st^ and 99^th^ quartile (whiskers). Scale bars are 10 µm throughout.

EGFP has a similar fluorescence lifetime of 2.09 ns and 2.04 ns when expressed alone or co-expressed with mCherry, respectively, however this is reduced to 1.73 ns if EGFP is fused directly with mCherry by a 12-amino acid linker (figure 2c-d). Forced proximity of mCherry therefore causes a shift away from zero of single pixel FRET efficiencies (figure 2e). We found that the integral of this curve between 0.1-1 provides a single metric describing the spread of single pixel FRET efficiencies. As changes in FRET efficiency are typically slight, it can be troublesome to determine their significance. We therefore present statistical comparison of these integral values as a non-biased means to conclude protein proximity.

FLIM-FRET analysis of EGFP-SNAP29 in the presence of mCherry-Stx17, demonstrates a strong reduction in fluorescence lifetime values when compared to donor alone (figure 2f-g), producing a FRET efficiency integral similar to fused EGFP-mCherry (figure 2h). The change in fluorescence lifetime (indicating molecular interaction) is restricted spatially to punctate structures, despite the presence of both proteins in the cytosol^7, 31^ and the evident SNARE-binding promiscuity of SNAP29^31^. This indicates that an unknown inhibitory mechanism likely prevents ectopic association.

To exclude the possibility that this interaction is the non-functional aggregation of over-expressed protein, a mutant Stx17 predicted not to form SNARE interactions was prepared by mutation of a conserved glutamine residue to glycine (mCherry-Stx17[Q196G]). This mutant demonstrated little deviation from donor-only data, reporting a FRET efficiency significantly smaller than for wild-type Stx17 (figure 2h), confirming a SNARE domain-mediated association of wild-type Stx17 and SNAP29. Mutant Stx17 does, however, still target and colocalise with SNAP29 (supplementary figure 6), highlighting the inadequacy of colocalisation for determining protein interactions.

FLIM-FRET analyses of EGFP-labelled endolysosomal R-SNAREs and mCherry-Stx17 report sub-8 nm proximity with EGFP-VAMP7, but not with EGFPVAMP8 (figure 3). As with Stx17 and SNAP29, expression of mutant Stx17[Q196G] abolishes FRET between EGFP-VAMP7 and mCherry-Stx17, highlighting the specificity of SNARE-domain association for our assay readout. We conclude VAMP7, not VAMP8 is the autophagosomal R-SNARE and that the regulated formation of a SNARE heterodimer acceptor on the autophagosomal membrane, analogous to other SNARE proteins^32^, is essential prior to VAMP7 engagement.

**Figure 3.**
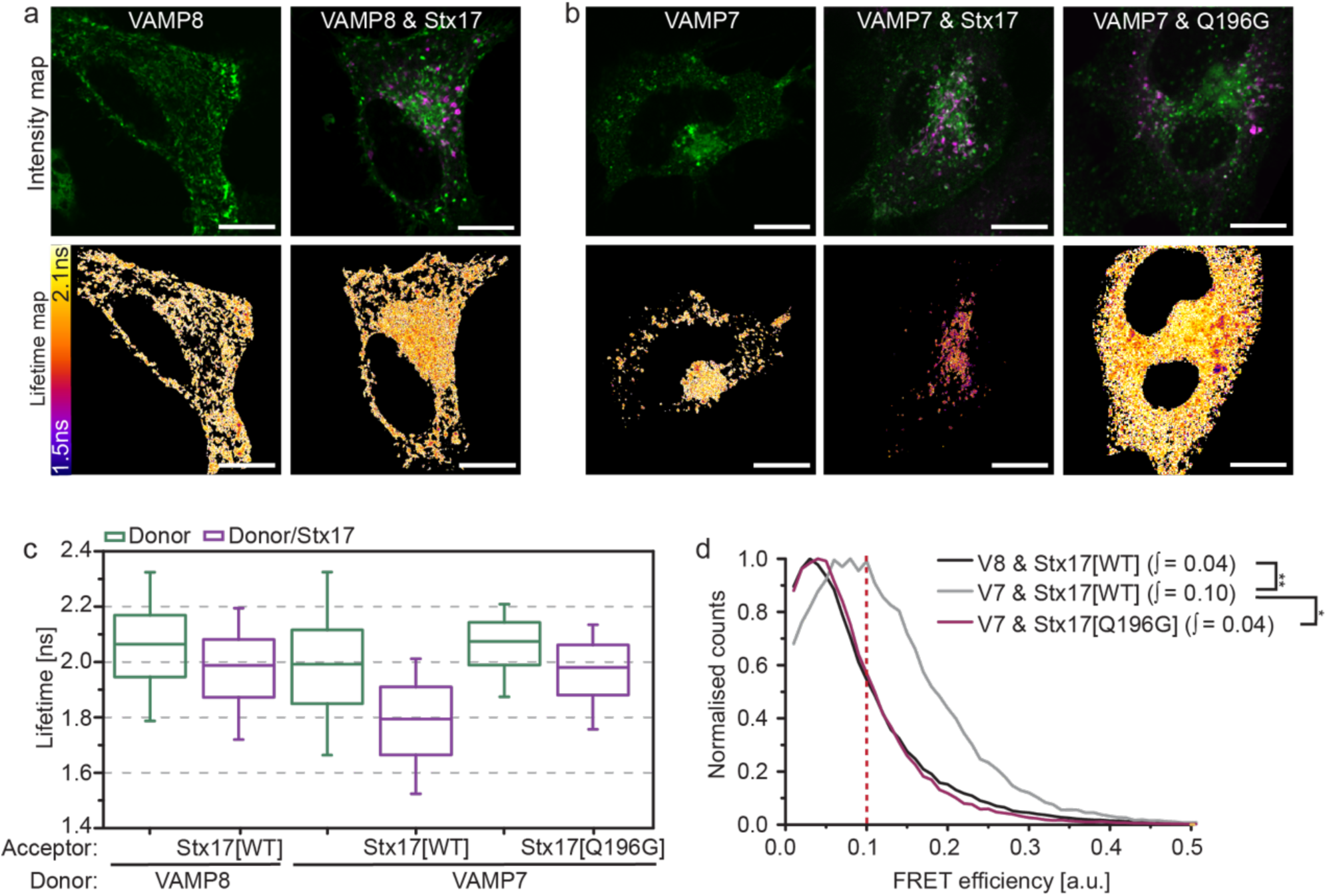
*In situ* FLIM-FRET identifies VAMP7 as a heterodimer-dependent interaction partner of Stx17. (**a-b**) Intensity images and their corresponding fluorescence lifetime maps probing changes in lifetime and proximity between donor only or donor and acceptor samples in rapamycin-treated cells, scale bars 10 µm. (**a**) EGFP-VAMP8 or co-expressed with mCherry-Stx17. (**b**) EGFP-VAMP7 or co-expressed with either mCherry-Stx17 or mCherry-Stx17[Q196G]. (**c**) Boxplots of the single pixel fluorescence lifetimes for the dataset presented in (a-b) and (**d**) the corresponding single-pixel FRET efficiency histograms; Integrated values above 0.1 were tested for statistical significance using a one-tailed unpaired two-sample t-test [n=4]. Box-and-whisker plots represent the median (central line), 25^th^ and 75^th^ quartile (box) and 1^st^ and 99^th^ quartile (whiskers). Scale bars are 10 µm.

### Autophagosomal SNARE interactions are modulated by Stx17 serine-2 phosphorylation

Given that SNARE proteins are highly reactive and promiscuous, we next addressed the question of spatial and temporal regulation of autophagic fusion. As Stx17 and SNAP29 colocalise, but do not interact, in the cytosol, this indicated to us that a regulatory step must exist to prevent ectopic heterodimer formation and uncontrolled fusion; the mode of regulation could be either inhibitory, or obligatory – i.e. the presence of an additional factor is essential to permit SNARE complex formation.

SNARE-mediated autophagosome clearance appears to be promoted at the fusion site by SNARE bundle-stabilising proteins, including HOPS-associated VPS33A^18^ and Atg14^33^. It remains unclear, however, if an inhibitory mechanism exists to prevent SNARE associations with Stx17 as suggested by our data in figure 2.

Syntaxin N-peptides can have a regulatory interaction with domain 1 of cognate SM proteins. However, the non-canonical sequence of this region in VPS33A led to the notion that it does not engage in this manner^22, 34^. Similarly, the Stx17 N-peptide is unusual in the syntaxin family in being highly charged; it consists of a string of four negative residues flanked by a predicted phosphoserine site with consensus for CK2^35^ (figure 4a-b). Stx17 serine-2 phosphorylation, which has been confirmed in phosphoproteome studies^36^, would have the effect of further increasing the net negative charge in this molecular region.

**Figure 4.**
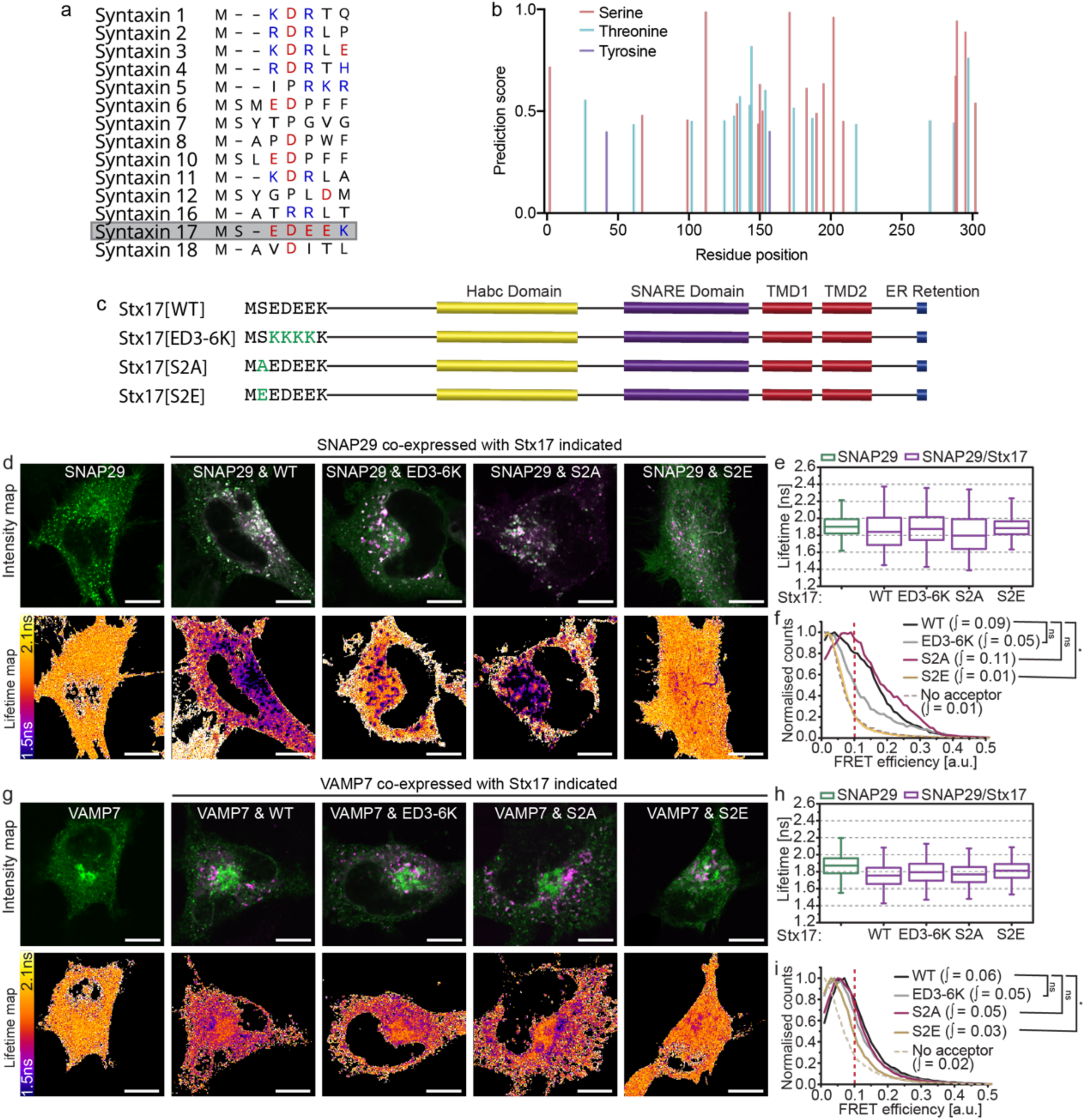
Stx17 N-terminal phosphoserine modulates SNARE complex formation. (**a**) An alignment of the N-peptide sequences of human syntaxin family proteins with negatively and positively charged residues indicated in red and blue respectively. Stx17, highlighted, demonstrates a highly negative patch. (**b**) NetPhos 3.1 phosphorylation site prediction scores for each residue of Stx17 indicates consensus for serine-2 phosphorylation. (**c**) Structure and sequence of wild-type Stx17 (Stx17[WT]) with designed N-peptide mutants: Stx17[ED3-6K] charge mutant, Stx17[S2A] phosphonull mutant and Stx17[S2E] phosphomimetic mutant. (**d**) Intensity and fluorescence lifetime maps of rapamycin-treated HeLa cells expressing EGFP-SNAP29 alone or alongside mCherry-fused Stx17[WT], Stx17[ED3-6K], Stx17[S2A] or Stx17[S2E] as indicated. (**e**) Single-pixel fluorescence lifetime boxplots and (**f**) normalised FRET efficiency histograms for the dataset presented in (d); four fields accumulated per condition. (**g**) Intensity and fluorescence lifetime maps of EGFPVAMP7 or co-expressed with Stx17 or one of its mutants as indicated in rapamycin-treated cells. (**h**) Single-pixel lifetime boxplots and (**i**) normalised FRET efficiency histograms for the dataset presented in (g); four fields accumulated per condition. FRET efficiency integral values above 0.1 were tested for statistical significance using a one-tailed unpaired two-sample t-test [n=4]. Box-and-whisker plots represent the median (central line), 25^th^ and 75^th^ quartile (box) and 1^st^ and 99^th^ quartile (whiskers). Scale bars are 10 µm.

We speculated that the negative N-peptide patch of Stx17 may enable a novel means of regulation that is modulated by serine-2 phosphorylation, similar to serine-14 phospho-regulation of syntaxin 1a^37^. To test this, we developed phoshonull, phosphomimetic and charge mutants: Stx17[S2A], Stx17[S2E] and Stx17[ED3-6K], respectively (figure 4c). No alteration in the localisation of any of the mutants was observed, meaning that their availability to binding partners was also unaffected. When FLIM-FRET was performed with these mutants to determine their interaction with either SNAP29 or VAMP7, we observed a profound, spatially restricted autophagosomal reduction in FRET only for the phosphomimetic mutant ([S2E]; figure 4d-i). No change was evident for the other mutants, indicating that increased negative charge at Stx17 serine-2 decreases SNARE complex formation, providing a putative inhibitory mechanism of autophagy regulation.

### Multi-modal regulation of Stx17 by VPS33A provides an autophagy master-switch

It is hard to conceive how a single charge mutation in the N-peptide of Stx17 alone could completely abolish detectable interaction with SNAP29 without the intervention of a modulating third party. Indeed, its N-peptide location hints at a second mode interaction with an SM protein, akin to syntaxin-SM proteins elsewhere in the cell^12, 41, 42%^. Importantly, however, an inhibitory role for VPS33A has been ruled out based on structural absence of an N-peptide binding pocket^33^ that appears to preclude interaction with monomeric syntaxin *in vitro*^22^.

It is known that the SM protein VPS33A promotes autophagosome clearance and its knockdown therefore blocks autophagic flux^18, 21^. Whilst the mechanism of action has never been established, the presence of a four alpha-helical binding groove in VPS33A^39^ hints that it can accommodate the SNARE bundle to physically stabilise cognate SNARE associations, as seen for other SNARE-SM pairings^12^.

To verify this, we recapitulated VPS33A knockdown studies to determine if its loss causes destabilisation of the autophagosomal SNARE bundle using a FLIM-FRET assay of ternary complex formation. Consistent with published work^18^, significant Stx17 puncta accumulation was observed in VPS33A siRNA-treated cells (figure 5a). This accumulation coincided with loss of the EGFPVAMP7 FRET observed in control knockdown cells when mCherry-Stx17 or mCherry-SNAP29 are present (figure 5b-d), validating for the first time *in situ* the proposed mechanism that VPS33A promotes autophagosome clearance by direct stabilisation of the SNARE bundle^18^.

**Figure 5.**
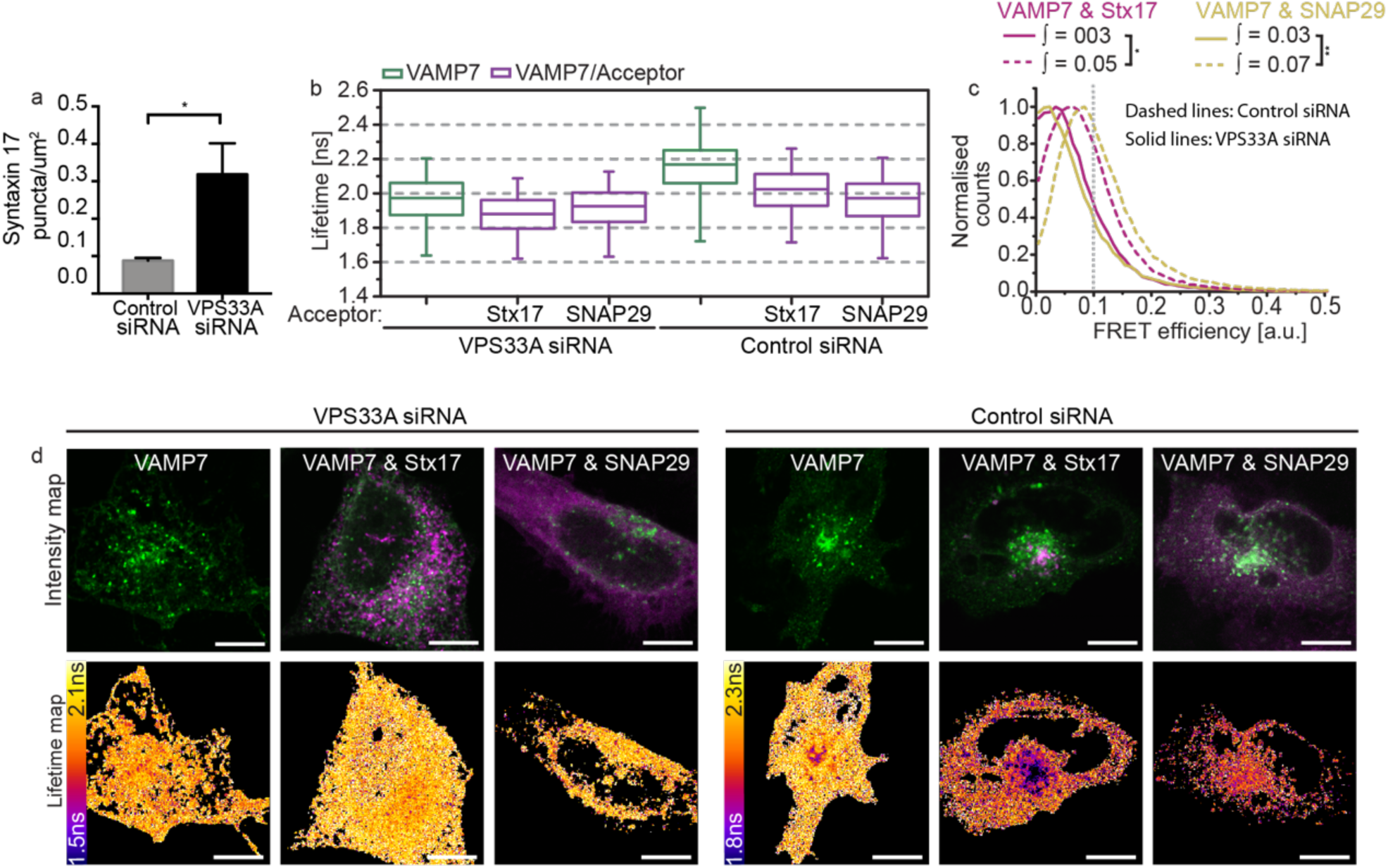
VPS33A promotes fusion by stabilising the SNARE bundle. (**a**) A barchart quantifying Stx17-positive puncta concentration per um^2^ in rapamycin-treated HeLa cells demonstrates accumulation of autophagosomes upon VPS33A knockdown (VPS33A siRNA) when compared to a non-targeting control knockdown (control siRNA). Significance tested using an unpaired two-sample t-test, n=4. (**b-d**) FLIM-FRET analysis of rapamycin-treated HeLa cells co-expressing EGFP-VAMP7 and mCherry-Stx17 or both mCherry-SNAP29 and non-tagged Stx17 when treated with either control siRNA or VPS33A siRNA. (**b**) Boxplots of all single pixel fluorescence lifetimes and (**c**) normalised single-pixel FRET efficiency histograms, both accumulated across four fields. (**d**) Representative intensity and fluorescence lifetime maps. FRET efficiency integral values above 0.1 were tested for statistical significance using a one-tailed unpaired two-sample t-test [n=4]. Box-and-whisker plots represent the median (central line), 25^th^ and 75^th^ quartile (box) and 1^st^ and 99^th^ quartile (whiskers). Scale bars are 10 µm.

To address our hypothesis that VPS33A may additionally bind monomeric Stx17, we carried out a structural analysis to assess binding potential. As VPS33A lacks the partially conserved N-peptide binding pocket of other SM proteins, we speculated that it may bind the Stx17 N-peptide through charge interactions. We observed strikingly positive electrostatic surface potentials of VPS33A position analogous to the N-peptide binding region in other family members^34^ (figure 6a), which we suspect interacts with the negatively charged Stx17 N-peptide in a manner mediated by serine-2 phosphorylation. Indeed, structural analyses of the HOPS complex have confirmed that both the syntaxin N-peptide binding region and the four alpha-helical binding groove of VPS33A are exposed when in complex^38-40^, which would therefore allow for multi-modal regulation.

To further define this interaction *in situ*, we temporally dissected the pathway by chemical inhibition of the fusion event. In the absence of baf A_1_, autophagic flux is unhindered and wild-type Stx17 demonstrates highly localised FRET with VPS33A in puncta-dense regions of the cell, reminiscent of the autolysosome (figure 6b-d). Surprisingly, this FRET is lost upon treatment with baf A_1_ to isolate pre-fusion vesicles, producing similar data as observed for the Stx17 phosphonull and charge mutants (figure 6e-g). This may indicate an accumulation of fusion-ready dephosphorylated wild-type Stx17, with altered VPS33A binding such that N-terminal proximities are no longer within the 8 nm range of FLIM-FRET sensitivity. In the case of Stx17[S2E], however, *increased* FRET efficiency was observed (figure 6e-g) compared to wild-type, with these interactions now localised to numerous cytosolic vesicular structures characteristic of autophagosomes. Notably, these differences in mutant Stx17 FRET are specific to VPS33A; Atg14, another promoter of autophagosomal SNARE assembly^33^, appears to interact with all Stx17 mutants (supplementary figure 7).

**Figure 6.**
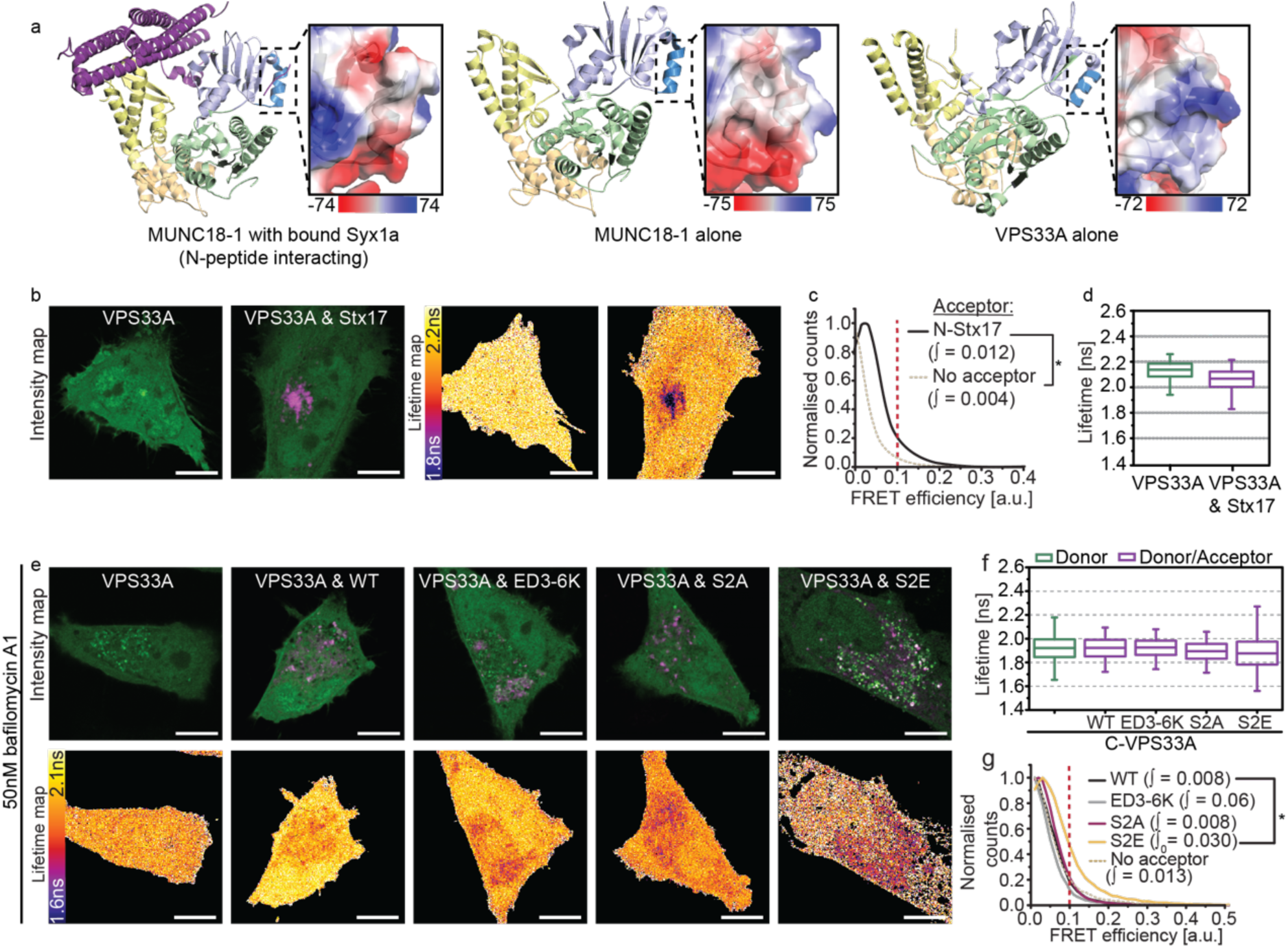
A phosphorylation master-switch in syntaxin 17 regulates autophagosomal SNARE assembly. (**a**) Cartoon diagrams of the structure of syntaxin 1a-bound Munc18-1 (left), Munc18-1 alone (centre) and VPS33A alone (right). SM protein structure is coloured by domain (domain 1 is light blue with dark blue highlighting the N-peptide binding region, domain 2 is green, domain 3a and 3b are yellow and wheat respectively). Magnified panels show the electrostatic surface charge in kT/e_c_ of the indicated syntaxin N-peptide binding region. (**b**) Intensity and fluorescence lifetime maps of rapamycin-treated HeLa cells expressing EGFP-VPS33A or co-expressed with mCherry-Stx17. (**c**) single-pixel normalised FRET efficiency histograms and (**d**) single-pixel fluorescence lifetime boxplots for the dataset presented in (b); accumulated over four fields per condition. (**e-f**) FLIM-FRET analysis of rapamycin- and bafilomycin A1-treated HeLa cells to probe for interactions between EGFP-VPS33A and mCherry-Stx17 on pre-fusion autophagosomes. (**e**) Intensity and fluorescence lifetime maps of VPS33A alone or in the presence of either wild-type Stx17 (WT) or its charge mutant (ED3-6K), phosphonull mutant (S2A) or phosphomimetic mutant (S2E). (**f**) Boxplots of all single pixel fluorescence lifetimes and (**c**) normalised single-pixel FRET efficiency histograms, for the datasets presented in (e); accumulated over four fields per condition. FRET efficiency integral values above 0.1 were tested for statistical significance using a one-tailed unpaired two-sample t-test [n=4]. Box-and-whisker plots represent the median (central line), 25^th^ and 75^th^ quartile (box) and 1^st^ and 99^th^ quartile (whiskers). Scale bars are 10 µm.

Taken together, these data suggest that VPS33A preferentially and transiently interacts with the phosphorylated Stx17 N-peptide prior to fusion. As VPS33A-Stx17 FRET follows an endolysosomal distribution in fusion-competent cells, it remains to be seen whether this is a pre-fusion population or if VPS33A re-associates with the Stx17 N-peptide post-fusion.

Thus, having begun to define the spatial organisation and regulation of the autophagic SNAREs, we sought also to place the VPS33A–Stx17 interaction mode(s) in temporal context within the autophagic fusion pathway by exploiting our Stx17 N-peptide mutants. First, we assessed the effect of specifically blocking autophagic flux using baf A_1_, an inhibitor of vacuolar H^+^ ATPase- and Ca-P60A/SERCA-dependent autophagosome-endolysosome fusion. Quantifying the density of LC3- or Stx17-positive puncta in cells expressing our Stx17 mutants thus dissects pre-fusion autophagosomes from the mix of pre- and post-fusion structures observed when autophagic flux is active (figure 7).

**Figure 7.**
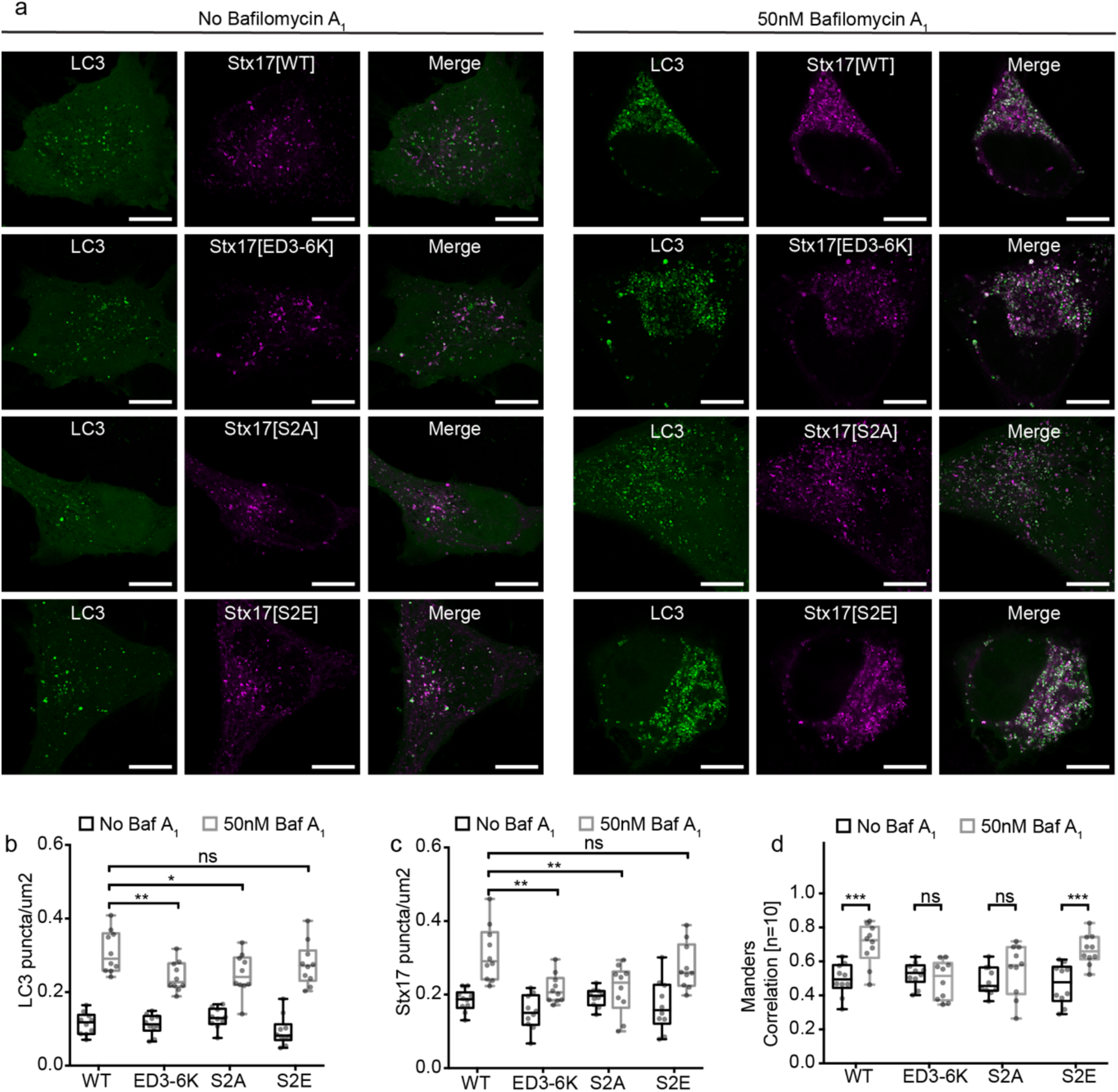
Stx17 N-peptide phosphorylation inhibits ectopic autophagosome clearance. **a**) Representative single channel and merged images of autophagic HeLa cells expressing LC3-EGFP and the variant of Stx17-mCherry indicated. Both fusion-competent cells (left panels) and fusion incompentent baf A_1_ treated cells (right panels) are displayed. Scale bars are 10 µm. (**b-c**) Boxplots of LC3-EGFP (c) or Stx17-mCherry (d) puncta concentration per cell in the presence or absence of baf A_1_. (**d**) Boxplots of Manders correlation coefficient to quantify colocalisation between channels in (a). Statistical significance tested with unpaired two-sample t-tests [n=10].

We confirmed that the autophagosomes containing each mutant were fusion competent by comparing LC3-EGFP puncta density in the presence or absence of baf A_1,_ finding a modest decrease in the post-treatment puncta in cells expressing Stx17[S2A] or Stx17[ED3-6K], and no difference in flux when Stx17[S2E] was expressed; all cells were fusion-competent (figure 7a and b).

Looking at the density of pre-fusion Stx17-puncta compared to non-treated cells revealed no effect of baf A_1_ on vesicle accumulation in cells expressing either Stx17[S2A] or Stx17[ED3-6K], compared to Stx17[WT] cells (Figure 7a and c). Conversely, we observed accumulation of LC3-positive puncta in Stx17[S2E] cells, that were indistinguishable from Stx17[WT]. Parallel analyses, looking at the colocation of Stx17 with LC3, to distinguish pre- and post-fusion were in agreement; baf A_1_ treatment resulted in a significant increase in LC3-Stx17 vesicles in Stx17[WT] and Stx17[S2E] cells, but interestingly, had no effect in Stx17[S2A] or Stx17[ED3-6K]-expressing samples. Taken together with our interaction data (figures 4-6), we contend that loss of this regulatory step in Stx17[S2A] and Stx17[ED3-6K] mutants permits ectopic clearance of autophagosomes by disengaging SNARE complex formation from SM protein action. Phosphorylation of Stx17 serine-2 is a prerequisite for VPS33A engagement, identifying Stx17 serine-2 as an essential spatial and temporal decision point for autophagosomal fusion to proceed.

## Discussion

We report that the autophagosomal Qa-SNARE, Stx17, functionally interacts *in situ* with the SNARE proteins SNAP29 and VAMP7 (figures 2 and 3), contrary to the current understanding that VAMP8 completes the heterotrimer. We have additionally begun to define the regulation of this interaction, demonstrating that VPS33A can either promote stabilisation of the SNARE complex or associate with monomeric Stx17 (figures 5-6), dependent on Stx17 serine-2 phosphorylation status (figure 4),which thus acts as an essential late checkpoint for fusion.

The spatial and temporal control of Stx17 interactions with SNAP29 and VAMP7 is essential to provide the exquisite regulation that must be applied to autophagy in the cell. Our data allowed us to propose a model of autophagosome clearance (overviewed in figure 8) where the availability of reactive Stx17 is modulated by a phosphosite on Stx17 serine-2. Pre-fusion, phosphorylation of this site contributes additional negative charge that prevents the formation of the Stx17-SNAP29 heterodimer acceptor for VAMP7, and so is an inhibitory step for autophagosome fusion with the endolysosome. This involves VPS33A association with the phosphorylated Stx17 N-peptide. However, alternate engagement of VPS33A with Stx17, likely *via* its four helical binding groove, has the effect of stabilising the SNARE bundle, enabling fusion of the opposing membranes. For this switch to occur, Stx17 serine-2 must be locally dephosphorylated by an as yet undefined phosphatase to release VPS33A domain 1 and our data suggest that this may happen transiently with the Stx17 N-peptide re-engaging with VPS33A post-fusion. It remains to be determined whether VPS33A actively inhibits phosphorylated Stx17, stabilises a self-inhibitory conformation of Stx17 or is required to recruit and transition Stx17 from an inactive to a reactive form.

**Figure 8.**
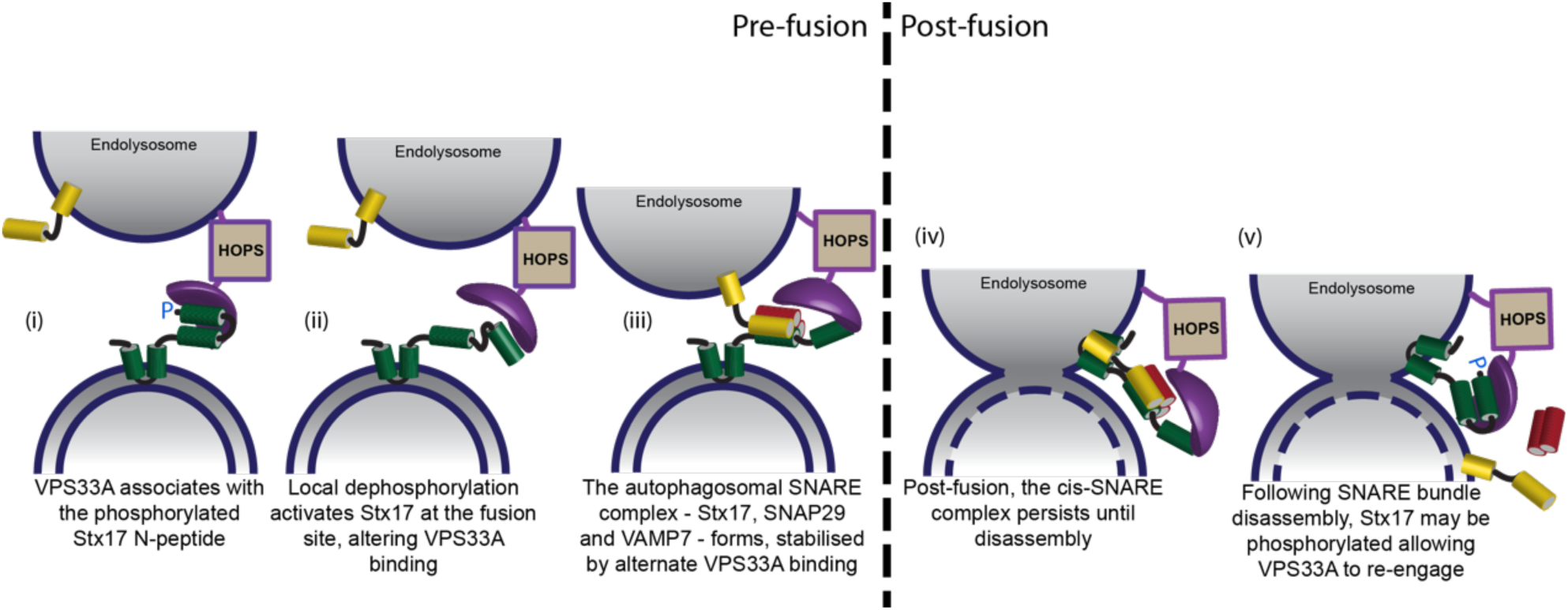
Proposed model of Syntaxin 17 interaction dynamics with VPS33A. Syntaxin 17 (green) and VPS33A (purple) interact via two distinct binding modes switched by the phosphorylation status of Stx17 serine-2. We propose that prior to fusion (i) VPS33A domain 1 associates with the phosphorylated Stx17 N-peptide, (ii) the Stx17 N-peptide is locally dephosphorylated at the fusion site, which alters VPS33A binding and enables SNARE bundle formation, providing a final stage regulatory mechanism, (iii) SNAP29 and VAMP7 may now associate with Stx17, forming a SNARE bundle that is stabilised by an alternate VPS33A association and which drives membrane fusion. Following fusion, (iv) the *cis*-SNARE bundle persists until actively disassembled, following which we suspect (v) Stx17 serine-2 is phosphorylated once more, allowing re-engagement of VPS33A domain 1 with the Stx17 N-peptide to deter further fusion events.

A central challenge in SNARE biology is to understand the interplay of the varied binding interactions of SM protein regulators. SM proteins can promote or inhibit SNARE-mediated fusion dependent on the binding mode^41, 42^, but the observed associations and their outcomes appear to vary for different syntaxin-SM pairings^34^. To reconcile these differences, it was recently proposed that two classes of SM proteins exist: class I, which have primary affinity for monomeric syntaxin and are characterised by N-peptide binding, and class II, which primarily bind the SNARE complex and lack N-peptide binding capability^22^. Our findings that VPS33A, the exemplar class II SM protein, has a structurally divergent but functionally analogous N-peptide binding patch, necessitates revision of this model.

Based on the data presented and their similarities with other mammalian syntaxin-SM interaction dynamics^43, 44^, we put forward a unified model of SM regulation where, *in situ*, N-peptide interactions are conserved across syntaxin-SM pairings, fulfilling a recruitment and tethering function. In the case of Syntaxin 1a, N-peptide tethering is sustained to aid binding mode transitions^45^, which, given the changes in FRET efficiency between phosphonull and phosphomimetic N-peptide mutants (figure 6), is likely accomplished by other means for Stx17. Indeed, VPS33A-associated HOPS components, VPS11 and VPS18, have been shown to associate with the Stx17 Habc domain^22^.

Our findings have additional implications for the regulation of autophagosomal clearance. Firstly, VAMP7 notably differs from VAMP8 in having a regulatory longin domain, consistent with the autophagosomal R-SNAREs identified in *D. melanogaster* and *S. cerevisiae*^28, 46^. This structural conservation may suggest an additional analogous regulatory mechanism that is as yet unexplored. Secondly, the evident inhibitory role of phosphorylation for autophagosome clearance may couple this process to intracellular energy levels based on ATP availability. If this is indeed the case, it would provide a fast feedback mechanism to prevent the unnecessary degradation of cellular material upon a return to nutrient replete conditions, bypassing the delayed onset of TORC1 reactivation^47^.

## Methods

### Cell culture, transfection and fixation

The HeLa cells used throughout this study were maintained at 37°C and 5% CO_2_ in Dulbecco’s modified Eagle’s medium supplemented with 100U/ml penicillin, 100ug/ml streptomycin, 10% heat-inactivated foetal bovine serum, 1X Glutamax and 1mM sodium pyruvate. Cells were cultured on poly-D-Lysine hydrobromide-coated glass coverslips and transfected using Turbofect Transfection Reagent (ThermoFisher Scientific) 24-30 hours prior to fixation with 4% paraformaldehyde and 0.1% glutaraldehyde. Samples were treated with 50mM ammonium chloride to reduce free aldehyde autofluorescence and mounted with Mowiol 4-88. In the case of knockdown experiments, siRNA was transfected using Lipofectamine RNAiMAX (ThermoFisher Scientific) 48 hours prior to DNA transfection.

### Plasmids

pEGFP-C1-Stx17, pmCherry-C1-Stx17 and pmCherry-N3-Stx17 were generated by ligating Stx17 isolated from pMRXIP-GFP-Stx17^7^ (Addgene, 45909) by PCR into the appropriate Clontech vector backbone. The mutant variants of pmCherry-C1-Stx17 (ED3-6K, S2A, S2E and Q196G) were obtained by site-directed mutagenesis using the QuikChange II Site-Directed Mutagenesis kit (Agilent). The same method was used to insert a stop codon at the C-terminus of Stx17 to generate Stx17[inv], a non-tagged variant of pEGFP-N3-Stx17. pEGFP- and pmCherry-C1-SNAP29, pEGFP- and pmCherry-C1-VPS33A and pEGFP-C1-VAMP8 were also generated by ligation into the appropriate vector of PCR-isolated SNAP29, VPS33A or VAMP8 from previously described plasmids^7, 18^ (Addgene, 45923, 67022 and 45919 respectively). pEGFP-C2-LC3^48^ (Addgene, 24920), pEGFP-C2-Atg14^49^ (Addgene, 21635) and pEGFP-C1-VAMP7^50^ (Addgene, 42316) were obtained from other groups and the former was used to generate pmCherry-C2-LC3 and pEGFP-mCherry-C2-LC3 by restriction and ligation. pEGFP-N3 and pmCherry-N1 (Clontech, discontinued) were used for FLIM-FRET control work along with pCDNA3.1-EGFP-mCherry, a fusion of two fluorescent proteins separated by a twelve amino acid linker, which was generated by isolation of the fusion protein from previously described pGEX-KG_EGFP-mCherry^45^ and ligation into the pCDNA3.1 vector backbone.

### siRNA

VPS33A knockdown was achieved using VPS33A stealth siRNA (ThermoFisher Scientific, HSS127975) with the following antisense sequence: 5′-AUGAGAUCUAAGCUGUACUCCUCCC-3′ Control knockdowns used siGENOME Non-Targeting siRNA #2, which is designed to target no known human genes (Dharamacon, D-001210-02).

### Autophagy assays

Autophagy was induced in cells by treatment with 160 nM rapamycin (ThermoFisher Scientific, PHZ1235) in normal growth media for one hour at 37°C and 5% CO_2_ prior to fixation. To assay for autophagic activity based on puncta accumulation upon inhibition of autophagosome-endolysosome fusion, cells were additionally treated where described with 50 nM bafilomycin A_1_ (Sigma-Aldrich, B1793) for 24 hours under normal culture conditions. Puncta accumulation was quantified per cell using the Spot Detector plugin^51^ within the open source software, ICY version 1.8.4.0^52^.

### Colocalisation image acquisition and quantification

Dual channel intensity images were acquired using a Leica SP5 SMD confocal laser scanning microscope (CLSM) fitted with a 63x 1.4 NA HCX PL Apo oil immersion objective lens. Samples were excited sequentially with 488 nm and 561 nm CW lasers and emission detected via internal photomultiplier tubes (Hamamatsu, R9624). Images were Nyquist sampled with a 1024 x 1024 pixel format. Images were processed by image deconvolution using a theoretical point-spread function (Huygens Software, SVI). Additionally, a rolling-ball background subtraction (Fiji) and thresholding to exclude non-punctate regions was carried out prior to Manders correlations. The Coloc 2 plugin (Fiji)^53^ was used for all colocalisations. Control colocalisation analyses were carried out by running the same algorithm on a fluorophore-dense 11 µm x 11 µm region of interest with a 90° transformation of the mCherry channel only.

### Stimulated emission depletion microscopy (STED)

Continuous wave (CW) gated-STED microscopy was performed with a 100X 1.4 NA HCX PL Apo oil immersion objective lens on a Leica SP5 SMD CLSM with STED capability. EGFP was excited using a white light laser tuned to 488 nm with an 80 MHz pulse rate and refinement of the point spread function was accomplished by concurrent depletion with an aligned CW 592 nm depletion laser. Emission wavelengths were isolated as for standard CLSM and detected with a time-gated Leica HyD hybrid detector synchronized to the excitation pulse. As the efficiency of depletion increases with depletion time^54^, time-gated detection between 0.5-12 ns was employed to ensure optimal depletion prior to detection. Images were acquired with a 0.015 µm pixel size to ensure Nyquist sampling.

### FLIM data acquisition

Fluorescence lifetime images were acquired on a Leica SP5 SMD confocal laser scanning microscope fitted with a time-correlated single photon counting module (PicoHarp 300) using a 63x 1.4 NA HCX PL Apo oil immersion objective lens. The donor EGFP was excited using a tunable white light supercontinuum laser operating at 488 nm and pulsing at 40 MHz. Emission was detected with an external single photon avalanche diode (MicroPhoton Devices). Recordings were integrated for 8-12 minutes to achieve maximum counts per pixel of approximately 10,000 photons, sufficient for accurate single pixel fitting (supplementary figure 8). Photon arrival times were recorded across 1600 time bins within the 25 ns window for each 161 nm pixel in a 256 x 256 format.

### FLIM-FRET data analysis

Single-pixel fluorescence lifetime analyses were carried out with SymPhoTime v5.4.4 (PicoQuant). A model fluorescence decay curve was generated from all pixels across a cell of interest and fitted bi-exponentially, as required for EGFP both in the absence and presence of mCherry (supplementary figure 8), using the maximum-likelihood estimation method. The resulting fit parameters were subsequently used to guide single pixel fitting. The output values for amplitude, lifetime and intensity were exported for graphical analysis and the generation of amplitude-weighted fluorescence lifetime maps, referred to in the text as ‘lifetime maps’, determined as

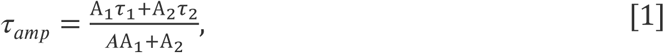

where A and τ are the amplitude and lifetime of the indicated exponent. To improve the reliability of the fit statistics, all pixels with less than 1,000 photon counts were excluded. FRET efficiency, was calculated as

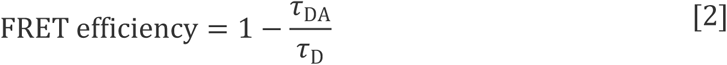

per pixel of the donor-acceptor image (τ_*DA*_) using the median amplitude-weighted fluorescence lifetime of all analysed donor-only pixels (τ_*D*_) per dataset.

## Acknowledgements

This work was funded by the Medical Research Council award (MR/K01563X/1) to RRD. We gratefully acknowledge the support offered by the MRC-funded Edinburgh Super-Resolution Imaging Consortium for both their expertise and infrastructure.

---

**Supplementary figure 1.**
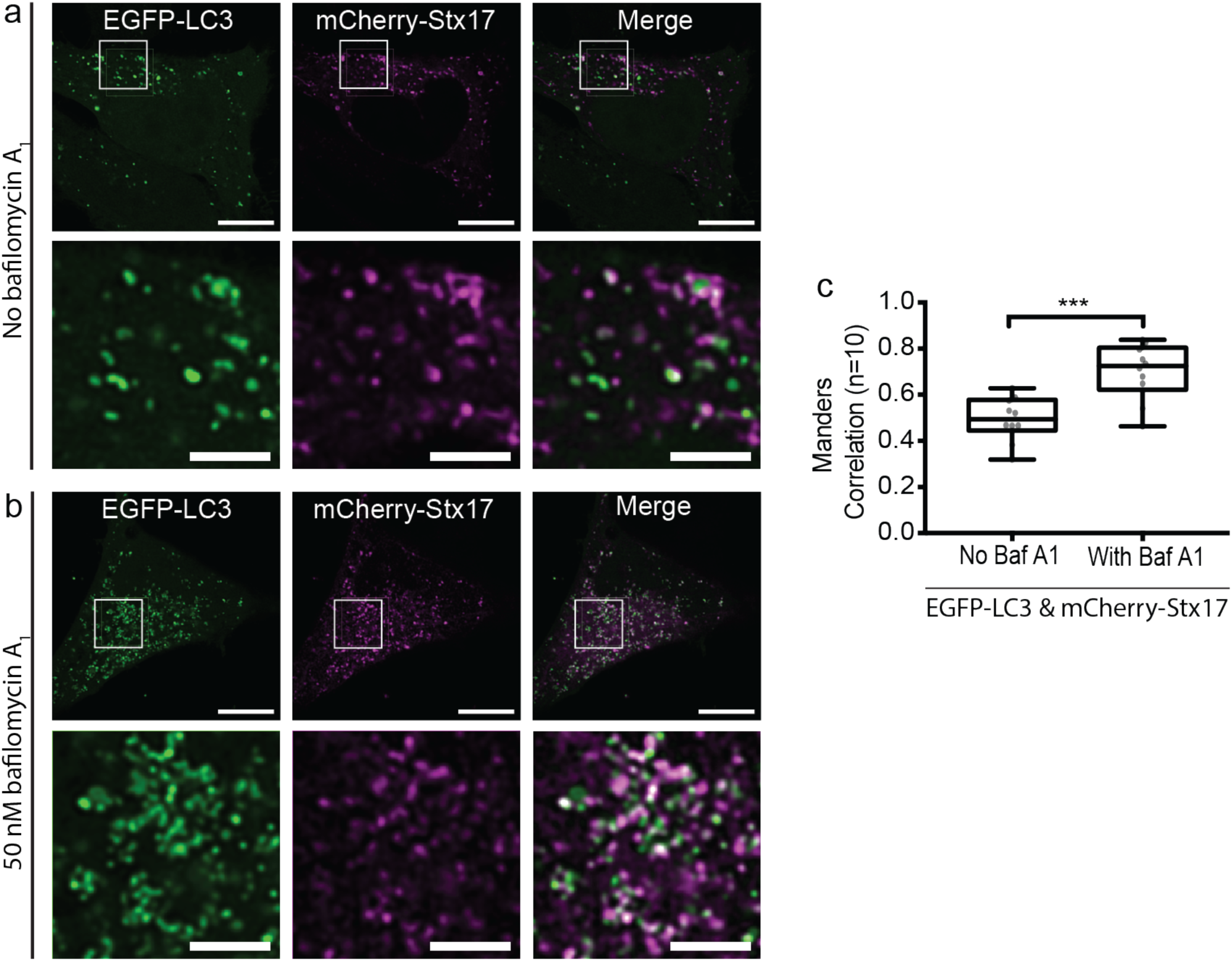
Stx17-EGFP resides on functional autophagosomes. (**a**) Single channel and merged CLSM images of autophagic HeLa cells co-expressing EGFP-LC3 and mCherry-Stx17 in the absence or (**b**) presence of bafilomycin A_1_. Scale bars are 10 µm for full field and 3 µm for enlarged images of indicated regions. (**c**) Box-and-whisker plots of Manders correlation coefficients to quantify data presented in (a) and (b). Statistical significance tested using an unpaired student t-test, showing a significant increase in correlation upon inhibition of fusion with bafilomycin A_1_.

**Supplementary figure 2.**
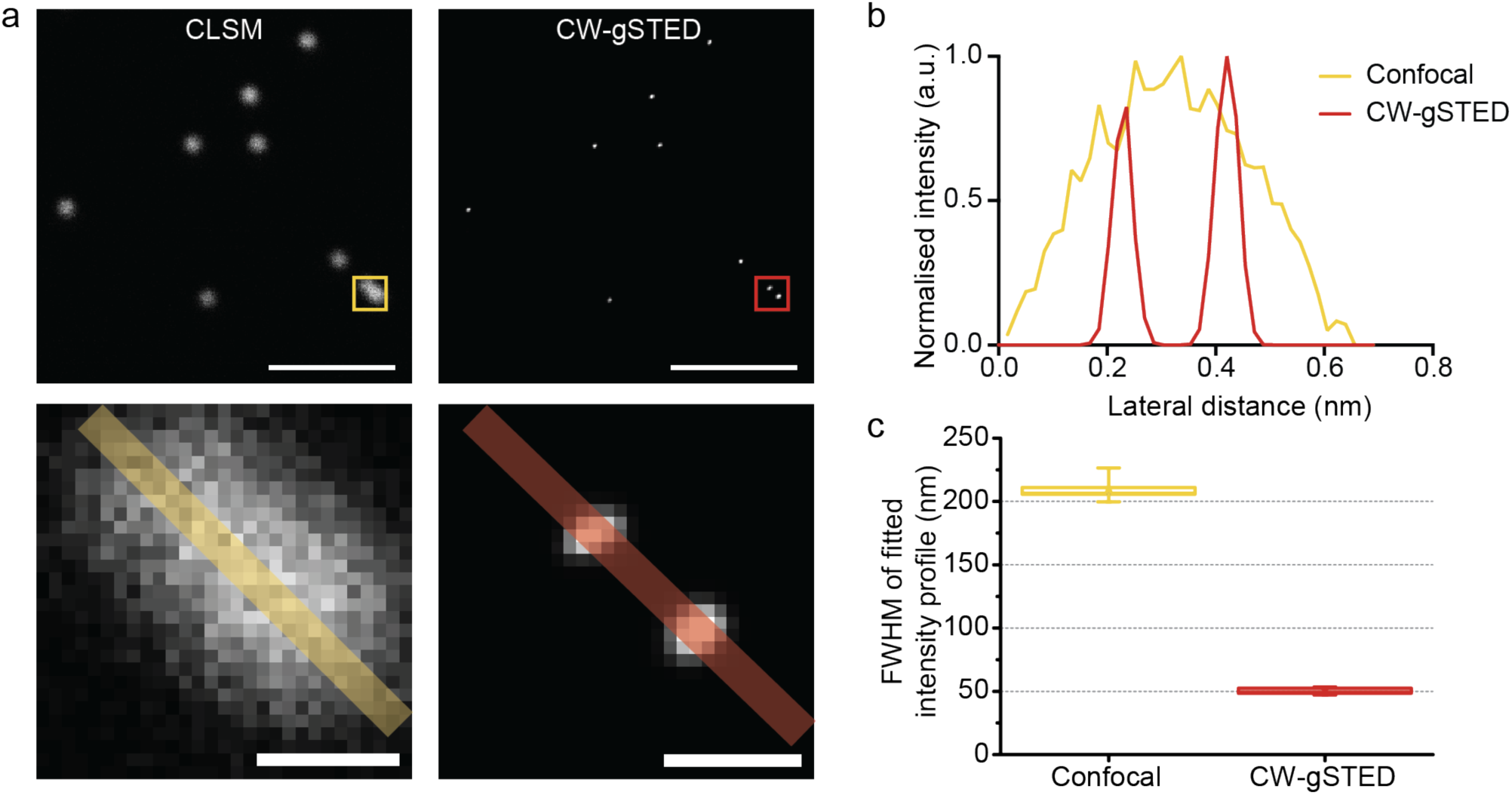
Achievable CW-gSTED resolution. (**a**) Confocal and deconvolved CW-gSTED images of 0.02 µm microspheres (505/515), with enlargement of boxed area below, scale bars are 10 µm (top) and 200 nm (below). (**b**) Intensity profile over the line indicated in (a) resolving two distinct peaks with CW-gSTED (red) and only one broad peak by confocal (yellow). (**c**) Box-and-whisker plots of the FWHM values through all single beads in (a) to quantify the improvement in resolution; the box represents 25^th^ to 75^th^ quartiles, the line represents the median value and the whiskers represent the maxima and minima.

**Supplementary figure 3.**
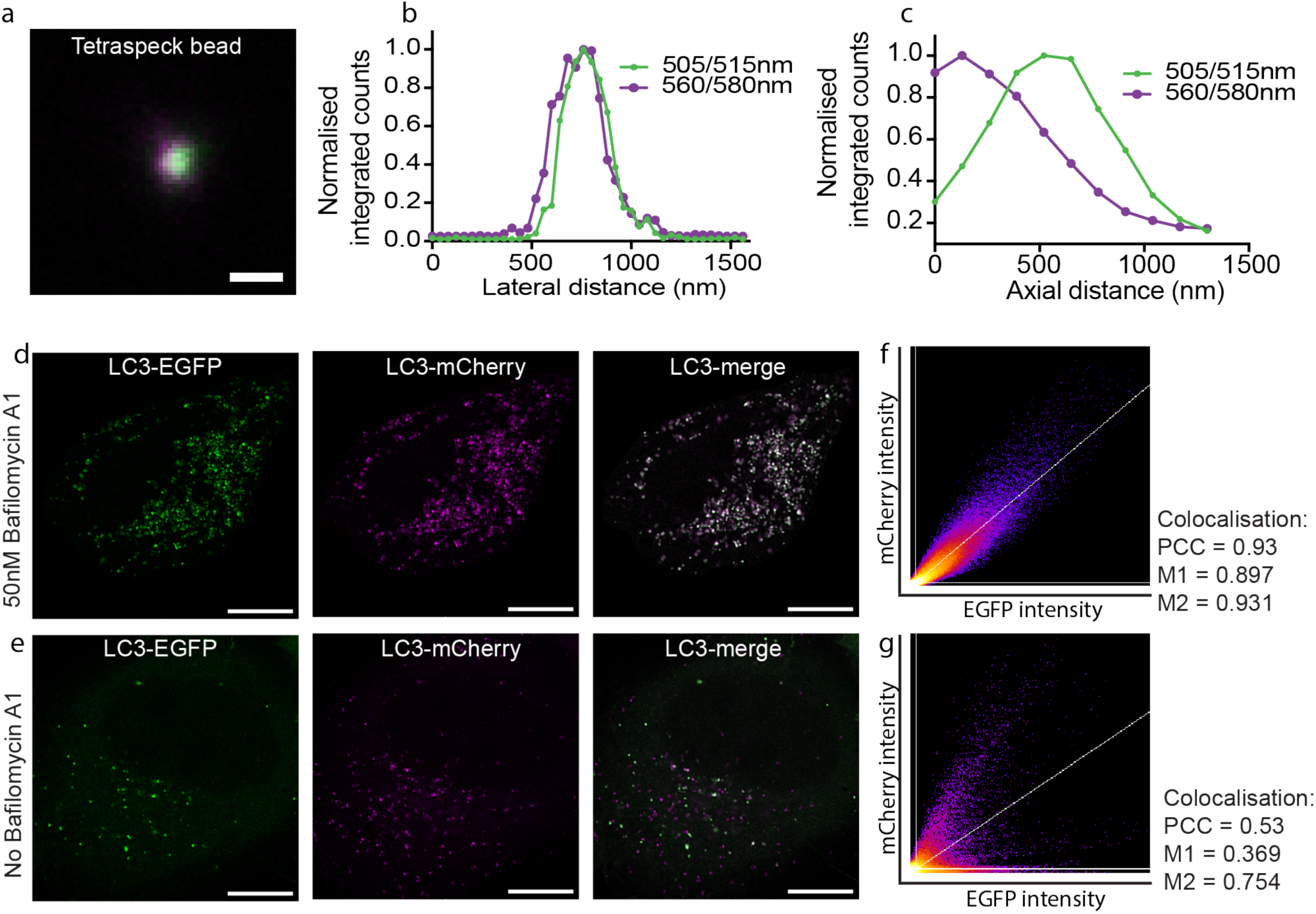
Fluorescence colocalisation acquisition and analysis. (**a**) A merged image of a single TetraSpeck bead with peak excitation and emission at 505 nm and 515 nm (green) or 560 nm and 580 nm (purple), scale bar 500 nm. (**b**) Lateral and (**c**) axial intensity profiles of the bead in (a). These indicate good lateral colocalisation but strong axial chromatic aberration, restrictinga colocalisation analyses to 2D datasets. (**d**) Intensity images of autophagic HeLa cells expressing dual-labelled LC3-EGFP-mCherry in the presence of bafilomycin A_1_ to prevent lysosomal degradation of EGFP and (**e**) in the absence of bafilomycin A_1_, scale bars are 10 µm. (**f**) A frequency scatter plot of the pixel intensities in (d), showing a linear relationship and (**g**) the same for (e), showing a non-linear relationship. As indicated, PCC reports high colocalisation for (d) but is more difficult to interpret for (e) where MCC demonstrates that as expected, only a subset of mCherry vesicles colocalise with EGFP (M1) while almost all EGFP vesicles colocalise with mCherry (M2).

**Supplementary figure 4.**
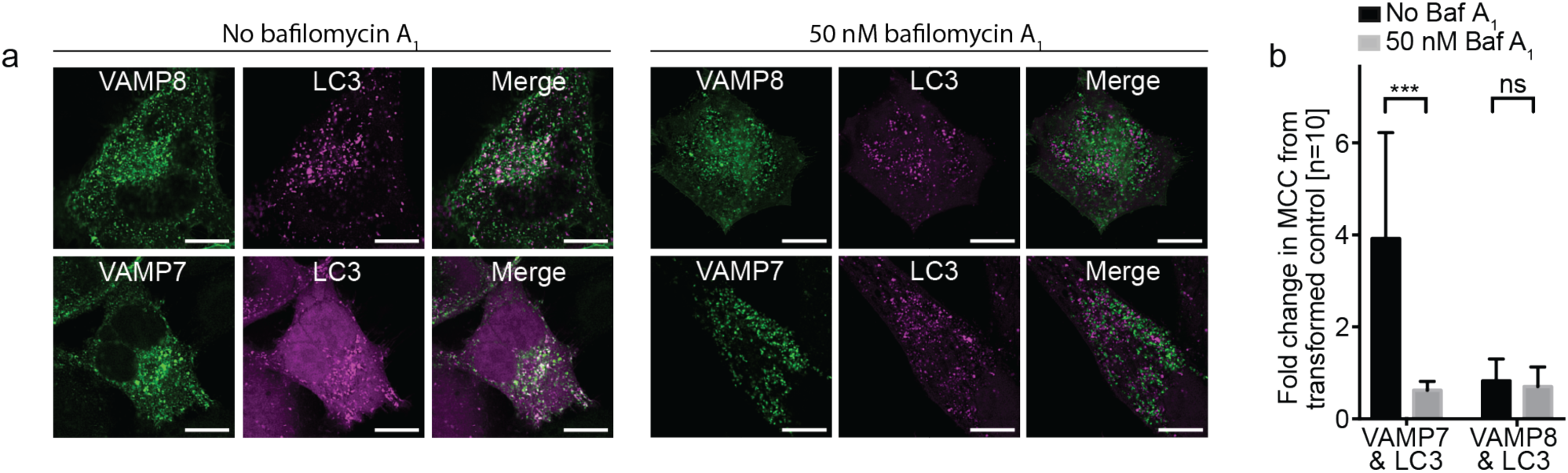
Fusion-dependent colocalisation of VAMP7 and LC3. (**a**) Single channel and merged representative CLSM images of autophagic HeLa cells co-expressing VAMP8 (top row) or VAMP7 (bottom row) with Stx17, both in the presence (right) or absence (left) of the autophagosome-endolysosome fusion inhibitor, bafilomycin A_1_. Scale bars are 10 µm. (**b**) A barchart comparing the fold increase in Manders correlation coefficients from the transformed negative control image. Statistical significance was tested with an unpaired sample t-test, showing a significant loss of correlation for VAMP7 and Stx17 when fusion is inhibited.

**Supplementary figure 5.**
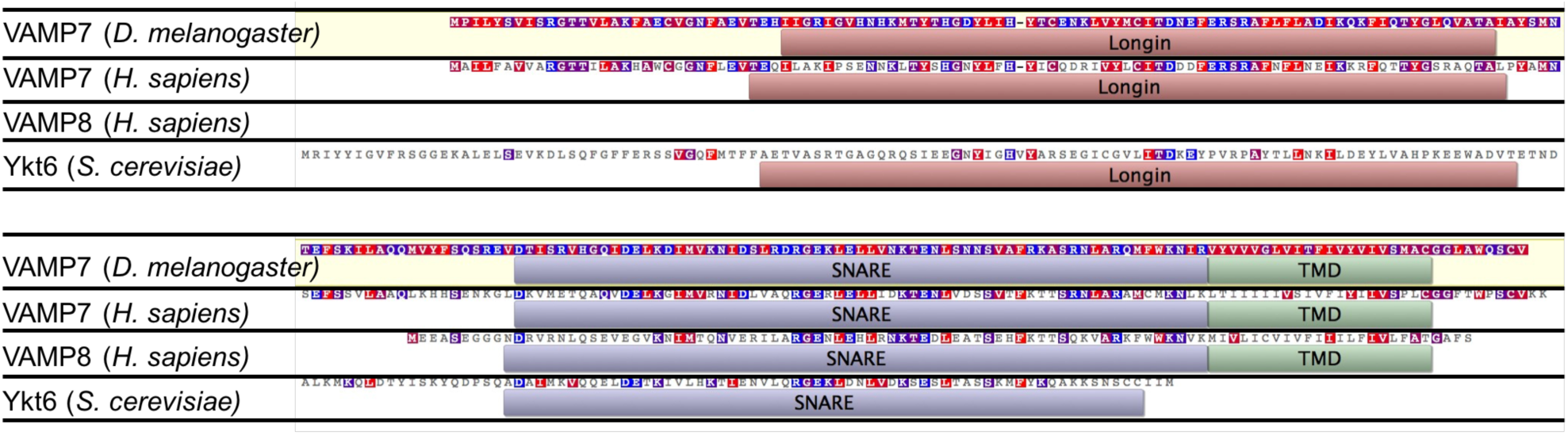
Putative autophagosomal R-SNARE alignment. Protein sequence alignments (wrapped over two lines) for the conflicting mammalian autophagosomal R-SNAREs, VAMP7 and VAMP8, in comparison to the identified autophagosomal R-SNAREs from other species, VAMP7 from *Drosophila melanogaster* and Ykt6 from *Sacchromyces cerevisiae*. Human VAMP8 alone lacks the longin domain that is conserved by all other putative autophagosomal R-SNAREs. Residues in consensus with VAMP7 from *D. melanogaster* (yellow) are coloured by hydrophobicity.

**Supplementary figure 6.**
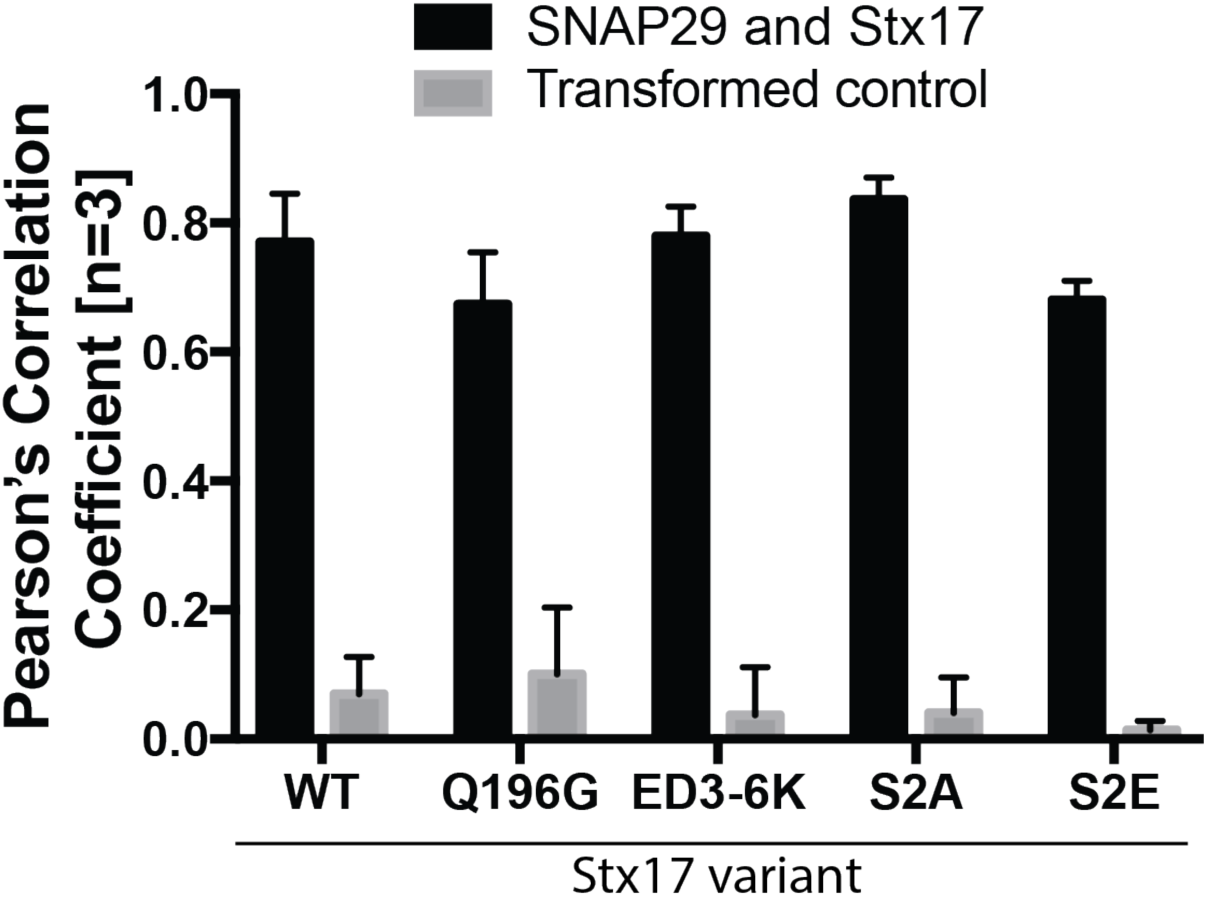
SNAP29 colocalises with mutant Stx17. A barchart of Pearson’s correlation coefficients to quantify the fluorescence colocalisation of EGFP-SNAP29 and wild-type mCherry-Stx17 (WT) or the various Stx17 mutants indicated. High correlation coefficients are evident for all variants of Stx17 despite the loss of interaction for both Stx17[Q196G] and Stx17[S2E], highlighting the limitations of fluorescence colocalisation for predicting protein interactions *in situ*.

**Supplementary figure 7.**
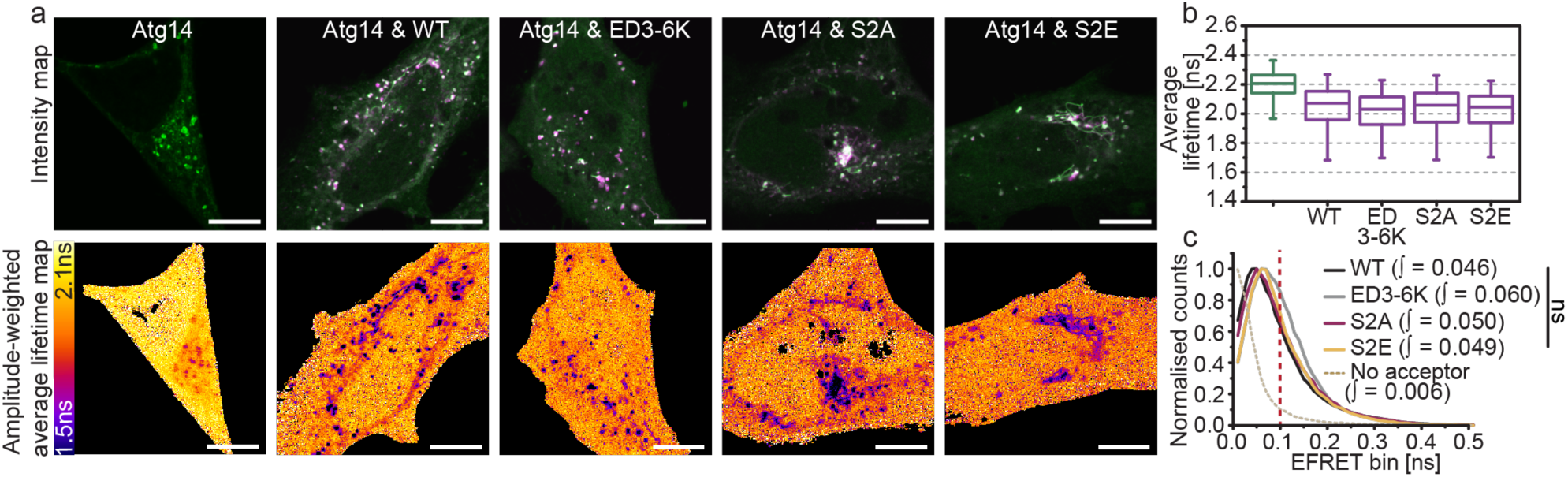
Atg14 interacts with wild-type Stx17 and its N-peptide mutants. (**a**) Intensity and fluorescence lifetime maps of rapamycin-treated HeLa cells expressing EGFP-Atg14 alone or alongside mCherry-fused Stx17[WT], Stx17[ED3-6K], Stx17[S2A] or Stx17[S2E] as indicated. (**b**) Single-pixel fluorescence lifetime boxplots and (**c**) normalised FRET efficiency histograms for the dataset presented in (a); four fields accumulated per condition. FRET efficiency integral values above 0.1 were tested for statistical significance using a one-tailed unpaired two-sample t-test [n=4]. Boxand-whisker plots represent the median (central line), 25^th^ and 75^th^ quartile (box) and 1^st^ and 99^th^ quartile (whiskers). Scale bars are 10 µm.

**Supplementary figure 8.**
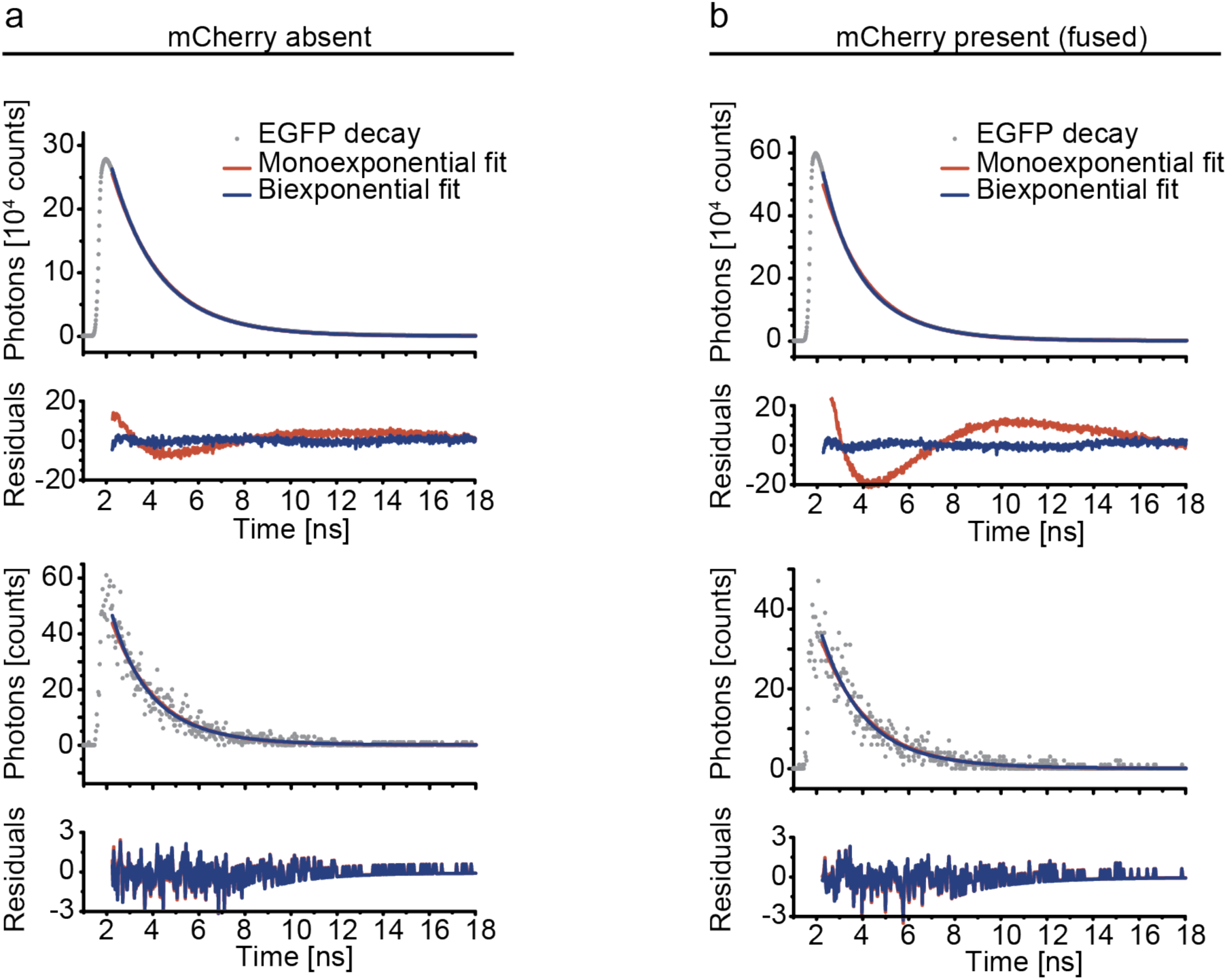
Representative EGFP fitted decays. (**a**) Whole cell (top) and single pixel (bottom) fluorescence decays for EGFP expressed alone in HeLa cells. Mono-exponential (red) and bi-exponential (blue) fits are shown with their corresponding weighted residuals below, indicating improved fit statistics with a bi-exponential equation. (**b**) As for (a) using data acquired from HeLa cells expressing EGFP fused to mCherry.

